# Identification and Establishing Floral calendar in, Jimma zone of Oromia Ethiopia

**DOI:** 10.1101/2023.09.12.557471

**Authors:** Tura Bareke, Kasim Roba, Admassu Addi

**Affiliations:** Holeta Bee Research Center, P.O. Box 22, Holeta, Ethiopia

**Keywords:** Floral calendar, Pollen analysis, abundance, monofloral honey, Honey flow period

## Abstract

Honeybee colony performance as well as production of honey and other hive products depends on bee forages. The study was designed with objective to identify and establish flowering Calendar of bee plants for multiple honey production. Bee forage inventory was made in a plot size of 20mx20, for woody plants and 2mx2m for herbaceous bee forages. Based on the inventory of bee forages, pollen load collection and honey pollen analysis, 103 plant species were identified as source of nectar and pollen for honeybees. From identified plant species Ageratum *conizoides, Guizotia scabra, Cordia africana, Datura arborea, Plantago lanceolatum, Rumex nervosus*, and *Justicia schimperina*, were the major honey source plants in the area.The fresh weight of pollen collected at different months of the year indicated that significant amount of pollen was collected during September, October to November but decline of incoming pollen weight was recorded during, March and July. The major honey flow period of the area occurred during November and extends early December and late February. The minor honey flow also resulted May-June. The pollen analysis of honey samples revealed that three monofloral honey types were identified which include Guizotia,Vernonia and Croton monofloral honey types comprising the pollen frequency ranging from 45% to 100%. The study concluded that floral calendar of bee forages strictly dependent on the seasonal availability of food sources and beekeepers should get awareness on management of honeybee colonies with the flowering calendar in the area to increase frequency of honey production.

## Introduction

Ethiopia has diverse climatic conditions and topography which favors for the growth of wide range of plants having different flowering pattern which result in availability of different honey flow periods which is entirely depend on flowering calendar of the area **[1]**.The floral calendar of bee plant is a timetable to indicate the date and duration of flowering of important bee plant species in an area. It is an important tool that informs the availability of certain bee forage for particular area, to predict time of honey flow period and their values to honeybees [2]. There are strong associations between floral calendar of bee plants and the seasonal cycles of honeybee colonies as it is affected by weather condition of the area [3]. The variations in rainfall and temperature can have influential effects on the onset & duration of the major bee plants, that can be coincide with management activities to increase yield of honey [4].

In the country the maximum flowering occurs after the main rainy season, followed immediately by dry season and small rainy season even though many species exhibit multiple flowering patterns within a year [5]. The length and intensity of rainfall influences the flowering of bee plants affecting the availability and quality of nectar and pollen. The well-defined sequences of colony events form a linear chain of relationship with rainfall: peak rainfall⟶peak flowering⟶maximum brood rearing⟶reproductive swarming [6] and [7].Therefore the timing of management operations corresponding to flowering pattern of bee plants which is critical in building up colony populations before the main nectar flow following the rainfall pattern of the area [8]. Even though honeybees naturally build up their population during periods when resources are available, the beekeeper must ensure that peak population is reached before or during the nectar flow. In Jimma zone, the area receives high rainfall for about 9 months as result, plants have extended flowering period and exhibit different flowering pattern [8]. As the result of this variation, beekeepers lack adequate information about appropriate flowering calendar of bee forages which is corresponding to seasonal colony management practices such as supering and honey harvesting.Therefore, identification of major bee forages and preparing floral calendar is important for determining appropriate honey flow period for multiple honey production. Thus study was designed to identify major bee forages and prepare flowering calendar and to assess the types of the mono floral honey produced in the area.

## Materials and Methods

### Description of the study area

The Jimma zone lies between 7^0^ 13’-8056N and 35° 52-37°37’E. The Zone is characterized by a humid tropical climate of heavy rainfall that ranges from 1200-2000mm. It has a relatively higher temperature of about 25°C- 30°C from January to April and having a minimum temperature of 7°C–12°C during the months of October– December.The zone also has relatively high forest accounting for about 56% of the country’s forest cover and different types of crops which include, coffee, maize, papaya, banana and many other root crops. The study districts includes Gomma, Tolay and Ommonada since they have a great potential for beekeeping.

## Methods

### Field observation

Plants visited by honeybees were identified through field observation. During observation the type of food source offered by plants, flowering periods and the behavior honeybees while collecting nectar and pollen were observed.

### Inventory of bee forages

To assess the bee forages abundance and diversity, four transect lines were laid out from apiary sites to North, South, West and East within 2Km radius following GPS assuming that honeybees efficiently utilize the resources within 2km radius. Apiary sites were selected systematically within 2 km distance from one to the other in order to avoid redundancy. Along these transects plots of 20mx20 m were laid out within 400 m interval between the sample plots. In order to retain accuracy, five (5) sub plots measuring 2mx2m (4m2) were laid out within the larger plot to capture herbaceous and grasses. All the plant species encountered in each sample plots were recorded and percentage cover of each species was estimated visually. For those plant species which could not identified in the field, sample of the specimens were collected using the standard Herbarium techniques and identified at National Herbarium of Addis Ababa University.

### Abundance of the species

The abundance of bee flora species defined here as the total number of all individuals’ species in all quadrate. Similarly the relative frequency of each bee flora species was calculated by determining the proportion of quadrate in which the species are encountered.

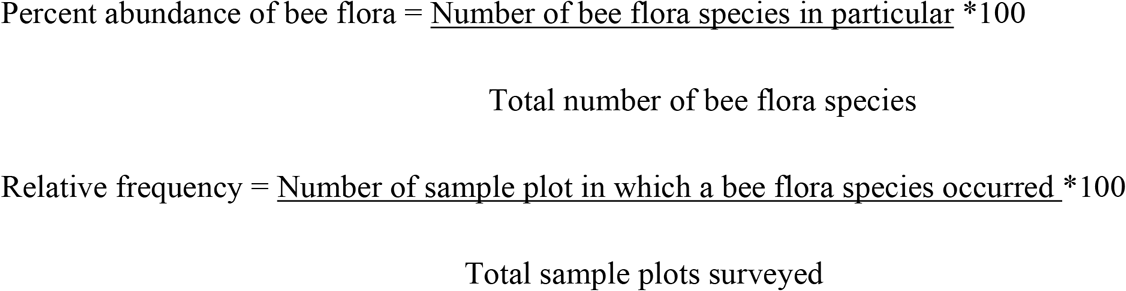

### Species diversity and Richness

To compare bee flora species composition among different districts, richness, Shannon diversity index, and Shannon evenness index was used, according to the following equation.

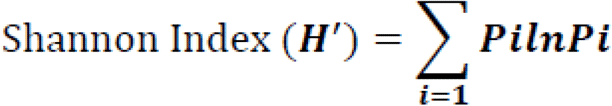

H’ is Shannon-Wiener diversity index, s is the number of species (pollen type), P _i_ is the proportion of individuals or the abundance of the ith species (pollen type), ln is the natural logarithm, and Σ is the sum from species 1 to species S. The evenness of the species was calculated based on the following Formula.

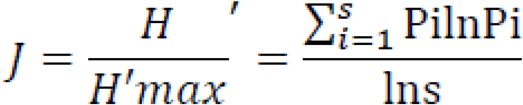

### Pollen load collection

Nine honeybees’ colonies were established at nine sites of the districts representing different agro-ecologies (lowland, mid land and highlands). For pollen collection honeybee’s colonies were fitted with pollen trap having 16% pollen trapping efficiency.The pollen grains were collected and sorted into colors and identified to the generic and species level.

### Honey pollen analysis

For honey pollen analysis, honey samples were collected from the different sites of the study area. The honey samples were analyzed for identifying species composition and frequency of pollen grains following the methods adopted by [9]. All types of the pollen grains were counted and scored as it occurred in the microscope field & percentage occurrence of pollen grains was used to determine their frequencies.

## Results

### Floristic composition

A total of 103 plant species belonging to 97 genera, and 78 families were identified in Jimma zone(Appendix1).TheFabaceae,Asteraceae,Solanaceae,Acanthaceae,Lamiaceae,Rutaceae, Moraceae,Myratceae,Acanthaceae, and Euphorbiaceae represented the highest number of species accounting for 22%, 17%.,10%,8%,8%, 6%, 6%,4%,7%, and 8% respectively. All remaining families consisted of species comprising less than 2% of having with low frequency occurrence in the area.

### Growth habit

The growth forms of bee forage utilized by honeybees comprises herb (35%), shrubs (32%), trees (29%) and climbers (4%) Fig. 2. The herbacous flora is dominant bee forages comprising weeds (*Guizotia scabra, Hypoestes forskaolii, Hypoestes triflora* and *Trifolium* spp) and some cultivated crops(*Brassica*,spp, *Vicia faba*, pulses cereals and oil crops also greatly contribute to honey production in the area.

**Fig 1.**
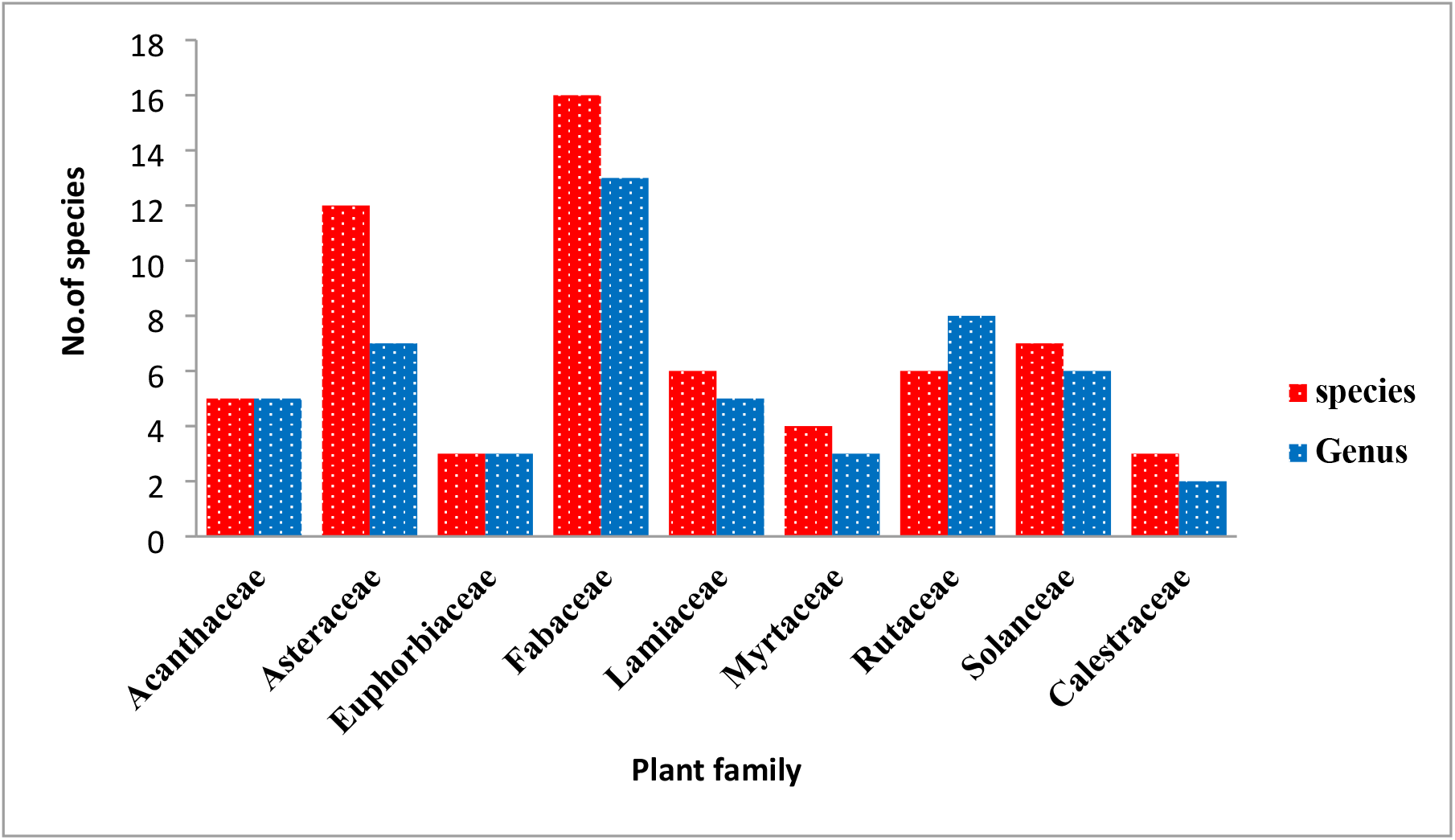
Percent of bee forage species composition and number of genera in west shoa zone.

**Figure 2.**
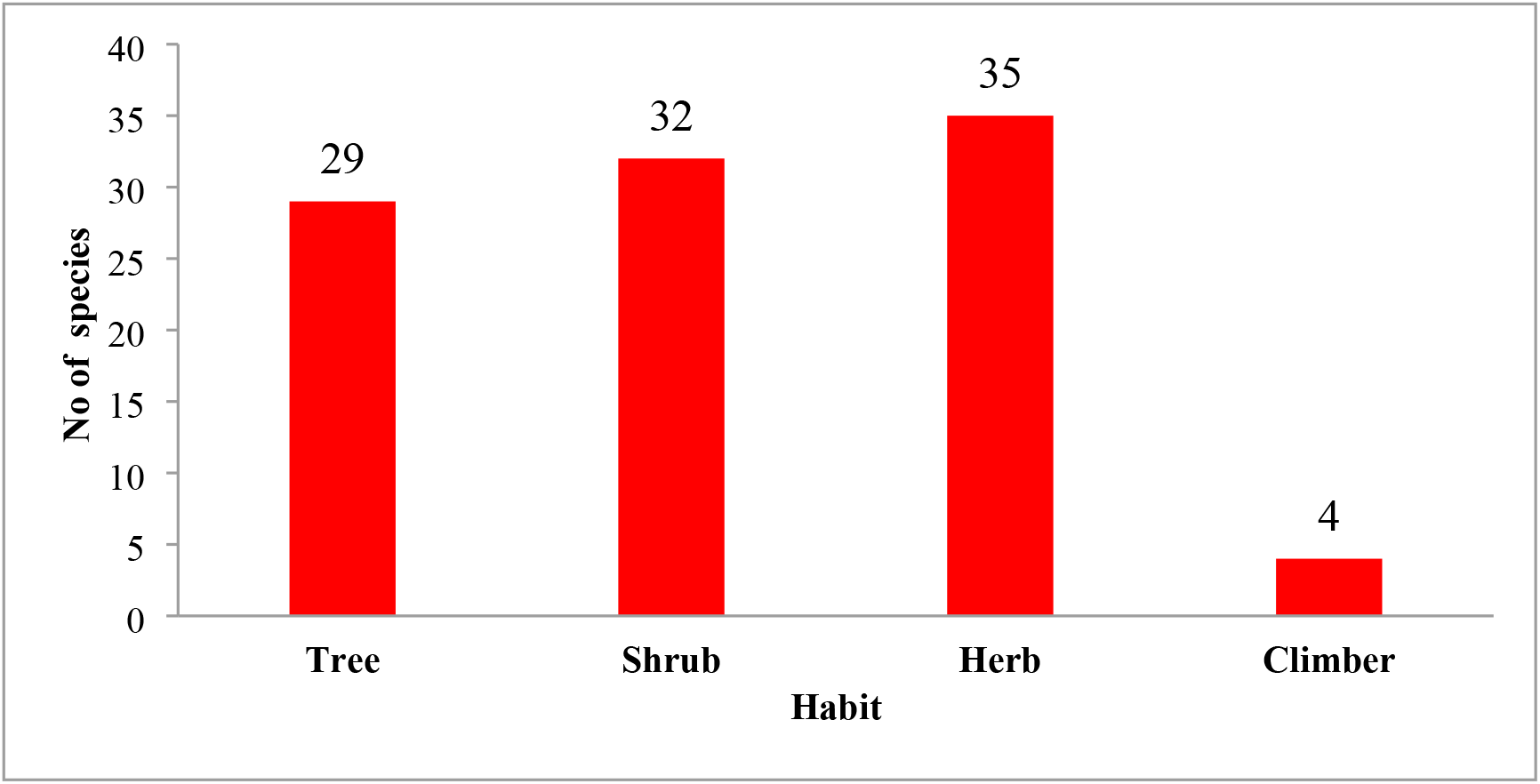
The growth habit of bee forages in the study area

**Figure 2.**
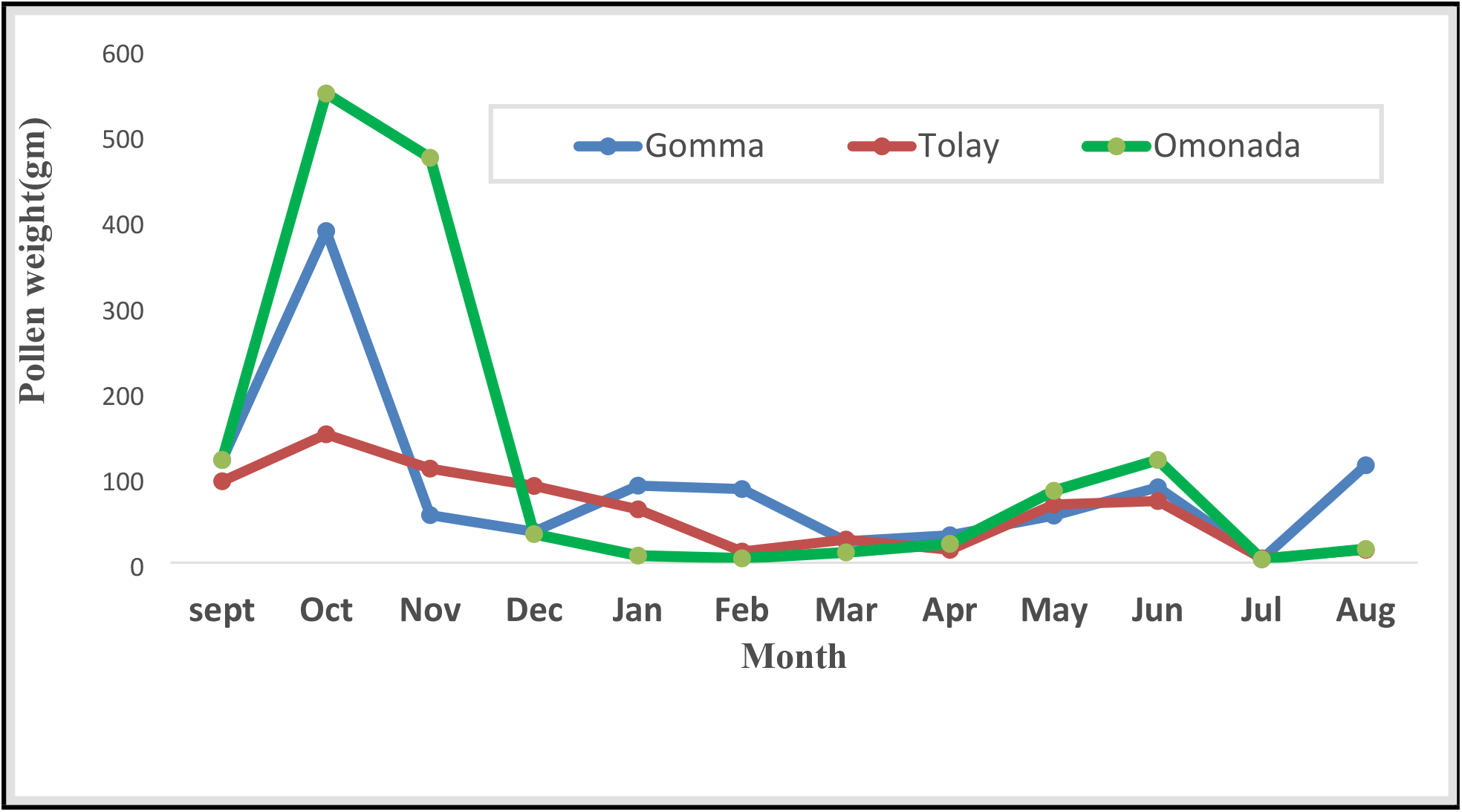
Monthly pollen collections in Jimma zone.

**Figure 3:**
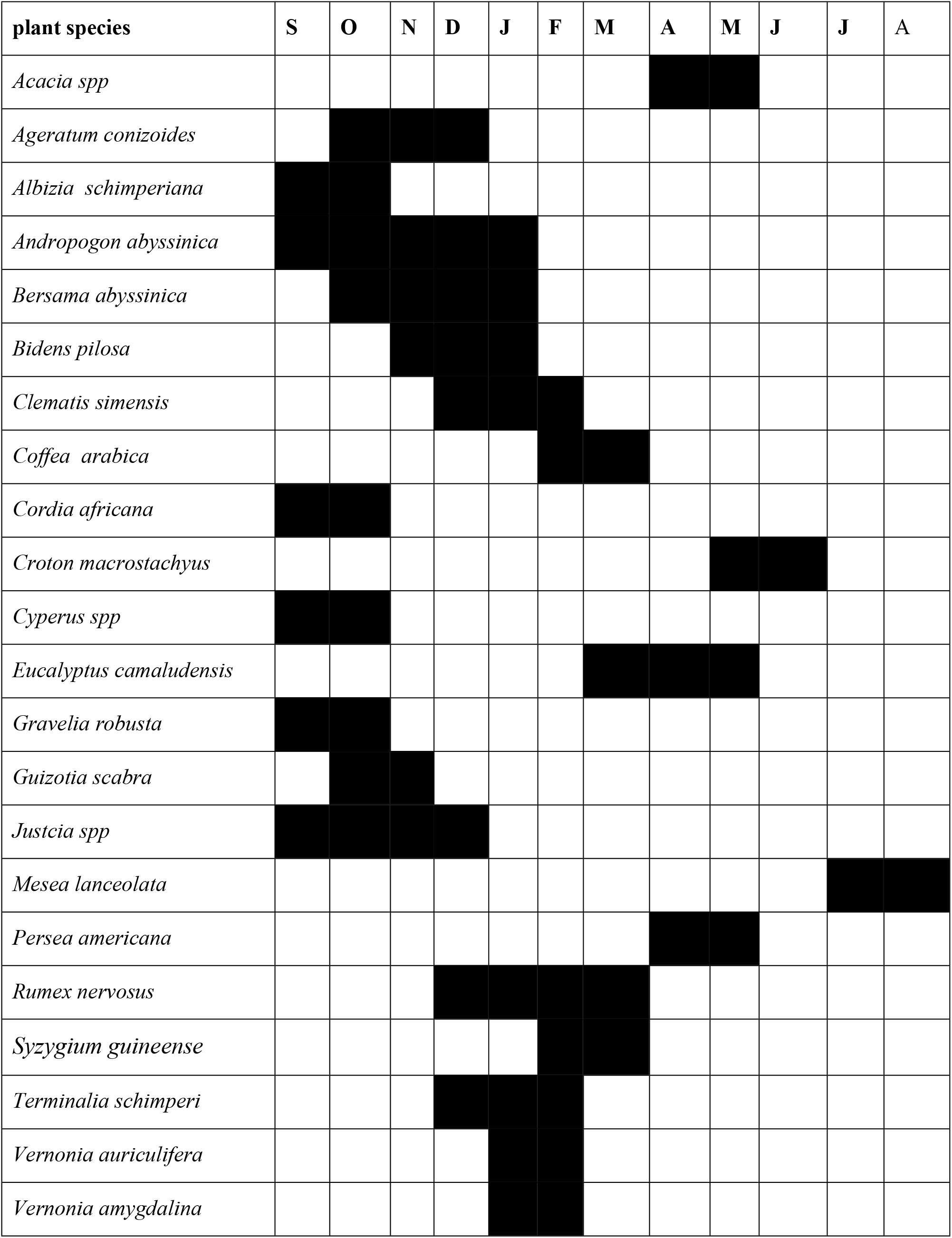
Flowering chart of major bee forages in Tolay, Gomma and Omonada districts.

### Abundance and diversity of bee flora

From field inventory of bee forages a few plant species are found to be the most frequent and abundant species in the area Table 1. These include *Eucalyptus* spp, *Croton macrostachysus, Justicia schimperana, Cordia africana, Vernonia* spp, *Guizotia scabra, Ageratum conyzoides, Achyranthes aspera* and *Hypoestes* spp and *Coffea arabica*.

**Table 1.**
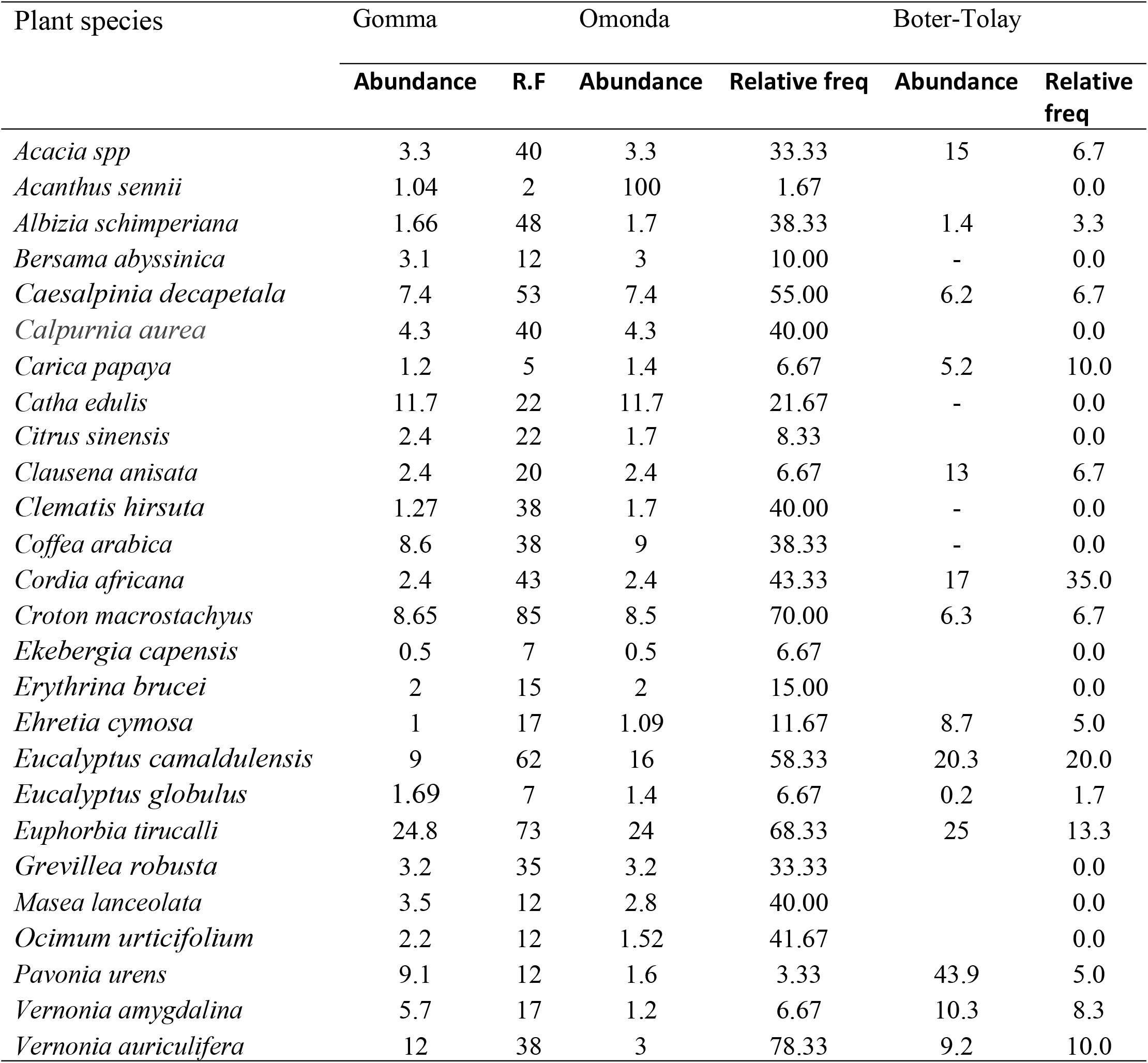
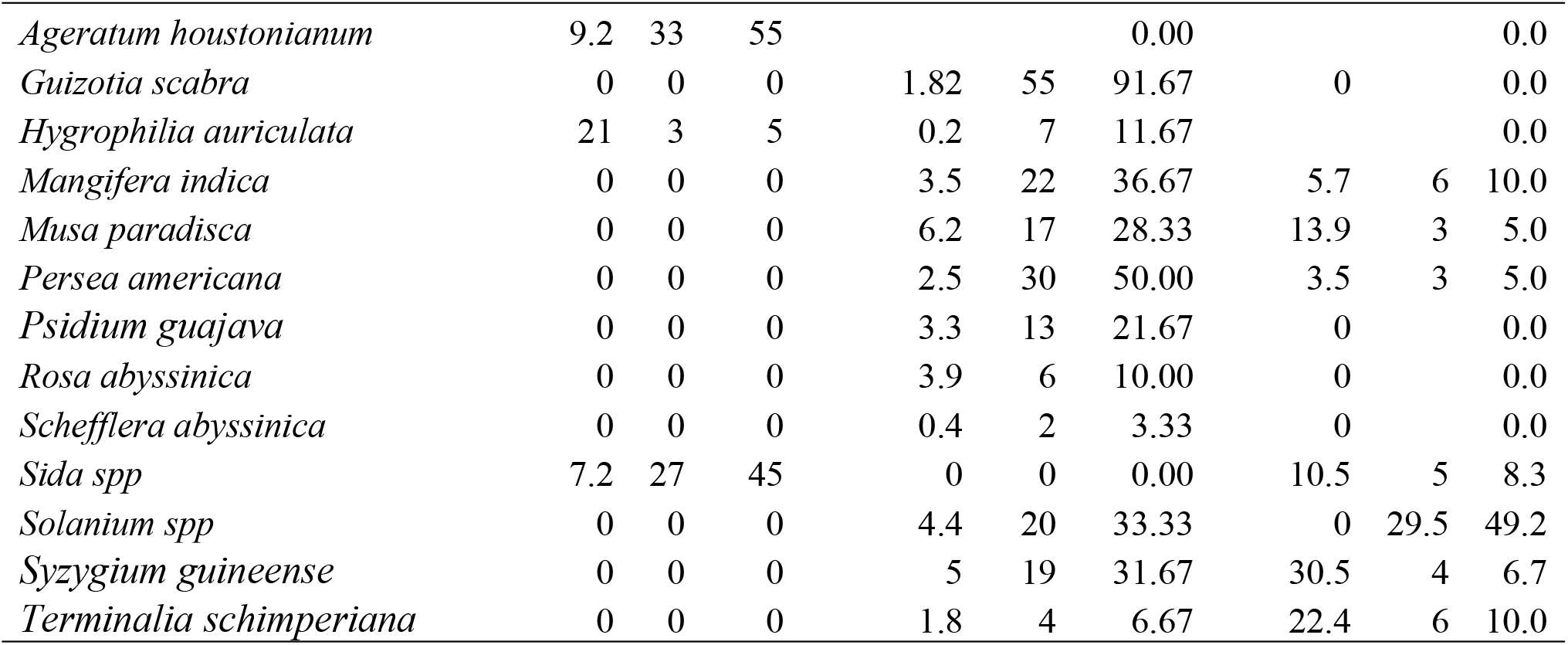
The abundance of major bee forages of the study districts.

| Plant species | Gomma |  | Omonda |  | Boter-Tolay |  |
| --- | --- | --- | --- | --- | --- | --- |
|  | Abundance | R.F | Abundance | Relative freq | Abundance | Relative freq |
| *Acacia spp* | 3.3 | 40 | 3.3 | 33.33 | 15 | 6.7 |
| *Acanthus sennii* | 1.04 | 2 | 100 | 1.67 |  | 0.0 |
| *Albizia schimperiana* | 1.66 | 48 | 1.7 | 38.33 | 1.4 | 3.3 |
| *Bersama abyssinica* | 3.1 | 12 | 3 | 10.00 | - | 0.0 |
| *Caesalpinia decapetala* | 7.4 | 53 | 7.4 | 55.00 | 6.2 | 6.7 |
| *Calpurnia aurea* | 4.3 | 40 | 4.3 | 40.00 |  | 0.0 |
| *Carica papaya* | 1.2 | 5 | 1.4 | 6.67 | 5.2 | 10.0 |
| *Catha edulis* | 11.7 | 22 | 11.7 | 21.67 | - | 0.0 |
| *Citrus sinensis* | 2.4 | 22 | 1.7 | 8.33 |  | 0.0 |
| *Clausena anisata* | 2.4 | 20 | 2.4 | 6.67 | 13 | 6.7 |
| *Clematis hirsuta* | 1.27 | 38 | 1.7 | 40.00 | - | 0.0 |
| *Coffea arabica* | 8.6 | 38 | 9 | 38.33 | - | 0.0 |
| *Cordia africana* | 2.4 | 43 | 2.4 | 43.33 | 17 | 35.0 |
| *Croton macrostachyus* | 8.65 | 85 | 8.5 | 70.00 | 6.3 | 6.7 |
| *Ekebergia capensis* | 0.5 | 7 | 0.5 | 6.67 |  | 0.0 |
| *Erythrina brucei* | 2 | 15 | 2 | 15.00 |  | 0.0 |
| *Ehretia cymosa* | 1 | 17 | 1.09 | 11.67 | 8.7 | 5.0 |
| *Eucalyptus camaldulensis* | 9 | 62 | 16 | 58.33 | 20.3 | 20.0 |
| *Eucalyptus globulus* | 1.69 | 7 | 1.4 | 6.67 | 0.2 | 1.7 |
| *Euphorbia tirucalli* | 24.8 | 73 | 24 | 68.33 | 25 | 13.3 |
| *Grevillea robusta* | 3.2 | 35 | 3.2 | 33.33 |  | 0.0 |
| *Masea lanceolata* | 3.5 | 12 | 2.8 | 40.00 |  | 0.0 |
| *Ocimum urticifolium* | 2.2 | 12 | 1.52 | 41.67 |  | 0.0 |
| *Pavonia urens* | 9.1 | 12 | 1.6 | 3.33 | 43.9 | 5.0 |
| *Vernonia amygdalina* | 5.7 | 17 | 1.2 | 6.67 | 10.3 | 8.3 |
| *Vernonia auriculifera* | 12 | 38 | 3 | 78.33 | 9.2 | 10.0 |

|  |  |  |  |  |  |  |  |  |  |
| --- | --- | --- | --- | --- | --- | --- | --- | --- | --- |
| *Ageratum houstonianum* | 9.2 | 33 | 55 |  |  | 0.00 |  |  | 0.0 |
| *Guizotia scabra* | 0 | 0 | 0 | 1.82 | 55 | 91.67 | 0 |  | 0.0 |
| *Hygrophilia auriculata* | 21 | 3 | 5 | 0.2 | 7 | 11.67 |  |  | 0.0 |
| *Mangifera indica* | 0 | 0 | 0 | 3.5 | 22 | 36.67 | 5.7 | 6 | 10.0 |
| *Musa paradisica* | 0 | 0 | 0 | 6.2 | 17 | 28.33 | 13.9 | 3 | 5.0 |
| *Persea americana* | 0 | 0 | 0 | 2.5 | 30 | 50.00 | 3.5 | 3 | 5.0 |
| *Psidium guajava* | 0 | 0 | 0 | 3.3 | 13 | 21.67 | 0 |  | 0.0 |
| *Rosa abyssinica* | 0 | 0 | 0 | 3.9 | 6 | 10.00 | 0 |  | 0.0 |
| *Schefflera abyssinica* | 0 | 0 | 0 | 0.4 | 2 | 3.33 | 0 |  | 0.0 |
| *Sida spp* | 7.2 | 27 | 45 | 0 | 0 | 0.00 | 10.5 | 5 | 8.3 |
| *Solanum spp* | 0 | 0 | 0 | 4.4 | 20 | 33.33 | 0 | 29.5 | 49.2 |
| *Syzygium guineense* | 0 | 0 | 0 | 5 | 19 | 31.67 | 30.5 | 4 | 6.7 |
| *Terminalia schimperiana* | 0 | 0 | 0 | 1.8 | 4 | 6.67 | 22.4 | 6 | 10.0 |

### Species richness and diversity

The analysis of Shannon diversity of bee forages indicated that there is variation of Shannon diversity index between three districts. The species richness also varied among the districts and’ relatively higher richness occurs in Gomma, and Omonada district as compared to Botertolay district Table 2.

**Table 2:**
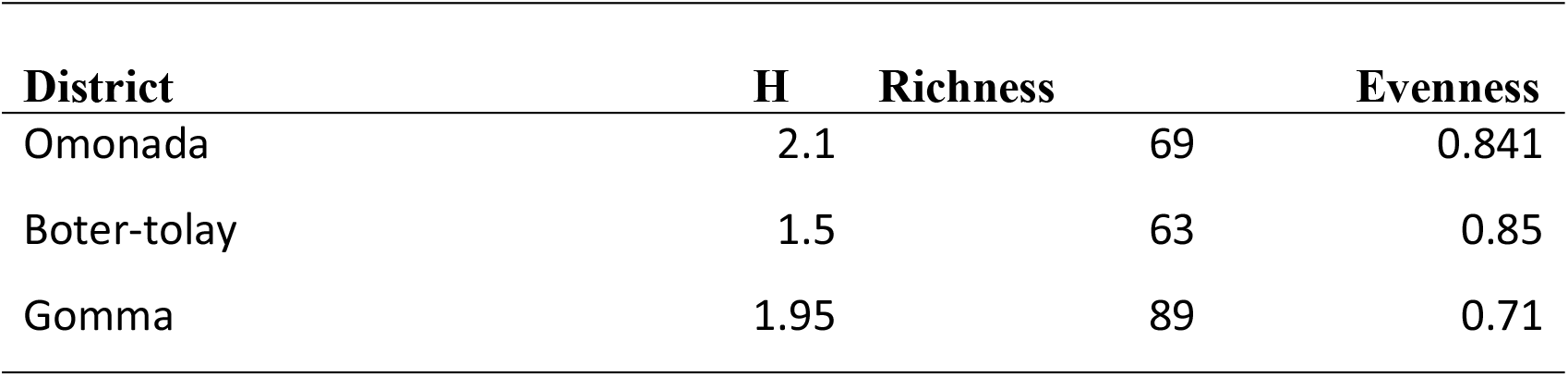
Species diversity of bee forages in west shoa zone.

| District | H | Richness | Evenness |
| --- | --- | --- | --- |
| Omonada | 2.1 | 69 | 0.841 |
| Boter-tolay | 1.5 | 63 | 0.85 |
| Gomma | 1.95 | 89 | 0.71 |

### Pollen load analysis of major bee forages

Among the identified pollen source plants, *Ageratum conizoides, Guizotia scabra, Cordia africana, Datura arborea, Plantago lanceolatum, Rumex nervosus*, and *Justicia schimperina*, were the most frequent pollen source plants and foraged for more than 50 days. The moderately foraged plant species were *Croton macrostachyus, Carica papaya, Gravillea robusta, Clematis simensis, Rhus vulgaris, Rosa abyssinica, Vicia faba* for foraged for 25-45 days and the rest of the species were least foraged by honeybees Table 3.

**Table 3:**
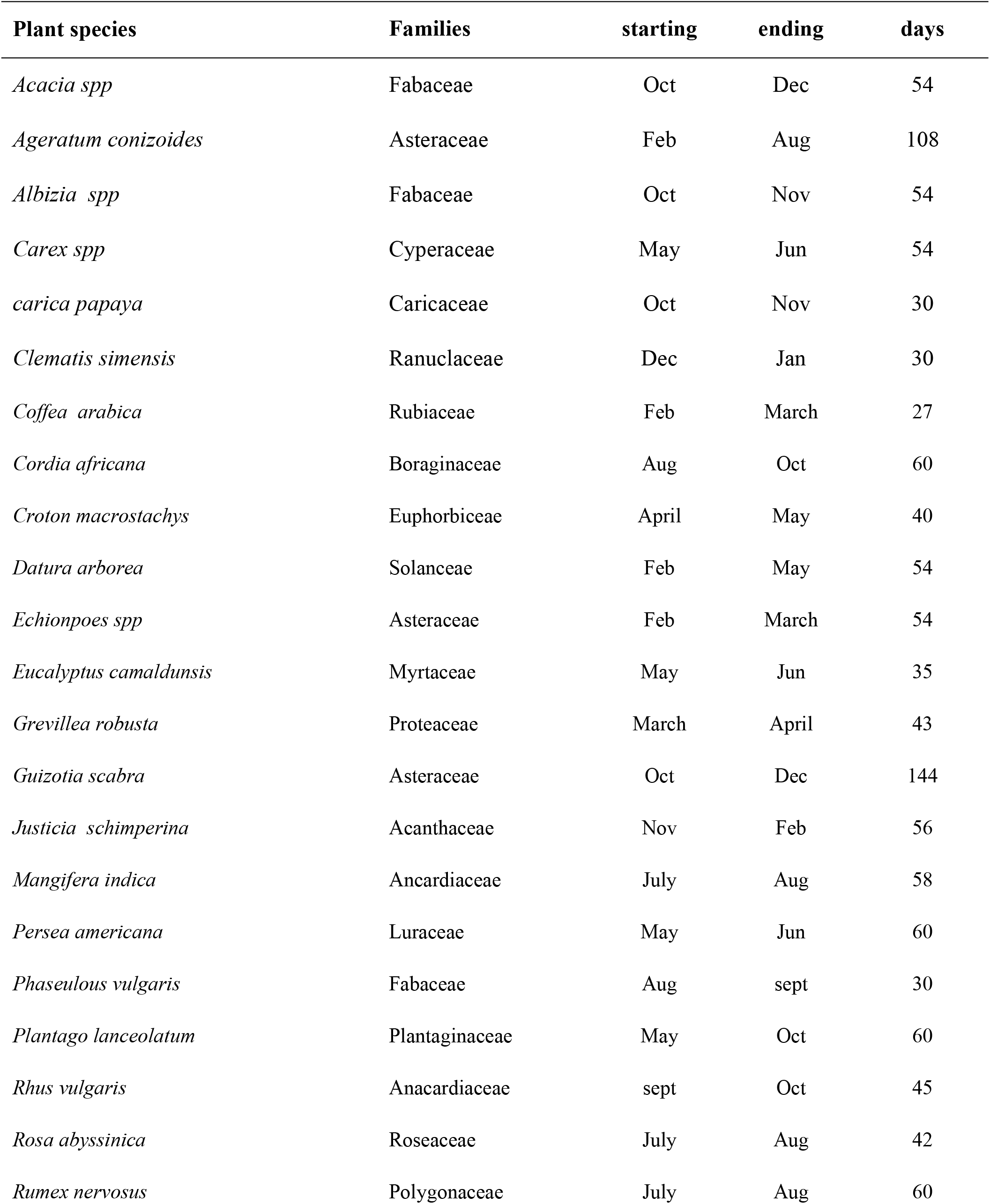
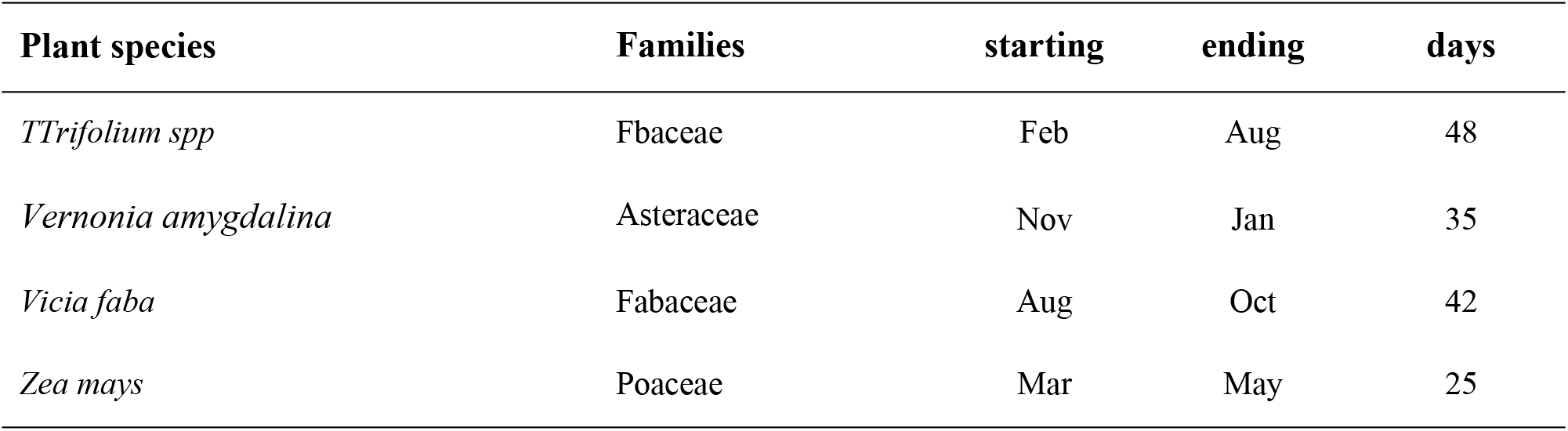
Foraging length of the major bee forages in Jimma zone.

### Seasonal availability pollen in the districts

Regarding the monthly pollen weight collection for Tolay, Gomma and Omonada districts, the highest pollen weight was recorded during the months of October, November and December. Relatively better amount of pollen was also collected during January and February in Gomma district. On top of this higher pollen weight was recorded during April to May in Gomma and Omonada as indicated in Figure **2**. The scarcity of pollen or very low pollen collection was recorded during February and July in Boter-Tolay and Omonda districts.

### Flowering calendar bee forages in area

The majority of bee plant species flowers in the area starting from mid-September to December reaching peak in November. About 30.6 % bee plant species flowers during spring (September to November) which includes *Ageratum conizoides, Andropogon abyssinica, Bidens* spp, *Cordia africana* and *Guizotia scabra*. Approximately 44.4 % of the plant species flowered during the dry season (December to February) including *Vernonia amygdalina, Coffea arabica, Bersama abyssinica* and *Clematis* spp. About 13.5% of the plant species flower during the small rainy season (March-May) which include *Eucalyptus camalduensis, Syzygium guineese, Mangifera indica* and *Croton macrostachyus* while 11.1% flowered during the big rainy season June-August (*Vicia faba, Zea mays and Maesa lanceolata)*. The analysis of flowering period of the bee forages, and seasonal pollen weight collection, three honey flow season were identified in Gomma district starting from mid-November to December from *Guizotia scabra* honey. The minor honey flow period was identified at end of January and March mainly from *Vernonia amygdalina* and *Coffea arabica*. On the other hand, *Croton macrostchyus* and *Eucalyptus* spp honey were harvested during months of May and June around Gomma, Omonada, and Tolay. The overall flowering period of major bee forages in Jimma zone was indicated in Figure 4.

**Figure 4.**
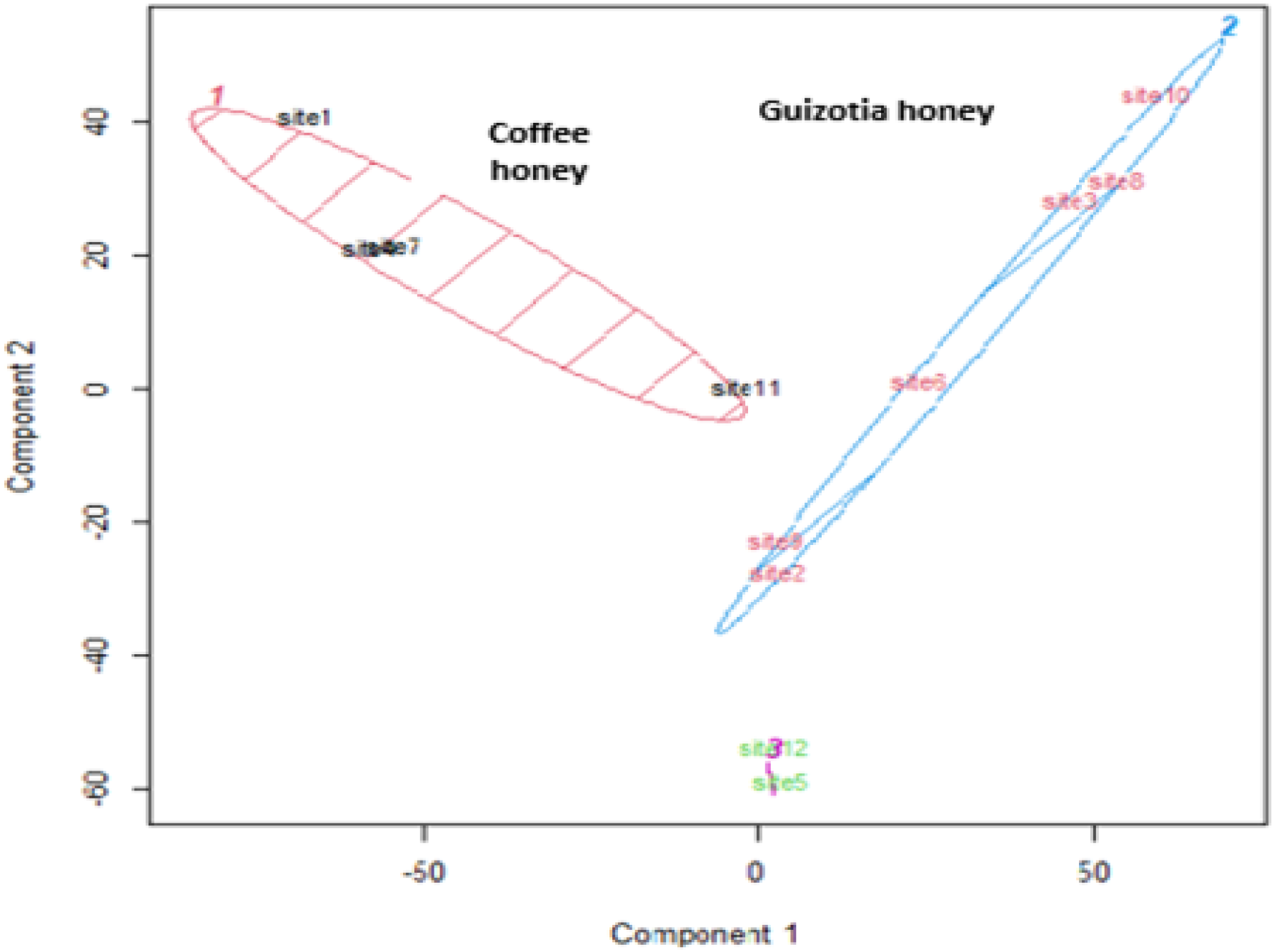
Monofloral honey types in Jimma zone

### Honey pollen analysis

From the pollen analysis of honey samples 39species representing 19 families were identified. The predominant pollen types (> 45%) identified was, *Guizotia scabra, Eucalyptus spp, vernonia* spp, *Croton macrostachys*; unidentified lamiaceae and *Coffee arabica (*Table 4). It is interesting to note that *Guizotia scabra* appeared as pre-predominant honey source plants during October honey flow season while *Croton macrostachys* and *Eucalyptus spp*, are dominant honey source plants during May–June and there is also honey from V*ernonia amygdalina, Coffee arbica* and *Albizia* spp were also reported during January and February. Finally, establishment of flowering calendar in the area shows that three honey flow seasons were identified of which September to October and April to May and during January and February.

**Table 4.**
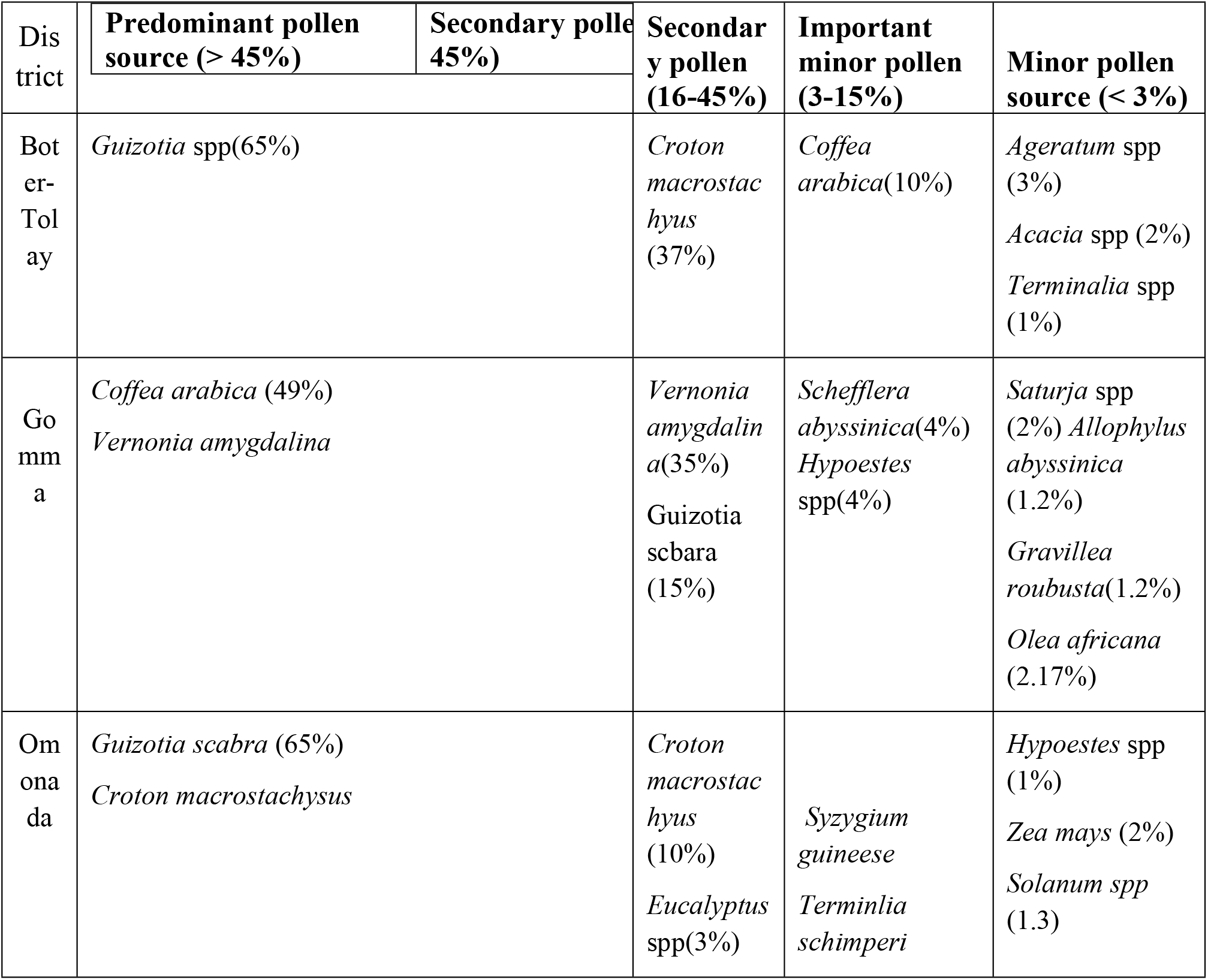
Predominate and secondary pollen source plants in west shoa zone.

### Monofloral honey

The classification of pollen percentage count using PCA clustering indicated that three monofloral honey types were identified, from Jimma zone. Accordingly, the Guizotia spp monofloral honey type covers large portion of cultivated areas of zone. On the other hand, *Coffea arabica* honey some degree of mixing with *Vernonia amygdalina* and *Croton macrostachyus* honey were found in Gomma district while *Croton macrostachyus* found in Omonada and Chora-Boter -Becho forest Figure 6. The pollen count percent dominance for monofloral honey ranges from 45 % to 100%.

## Discussions

From pollen load collection and honey pollen analysis,103 plant species were identified belonging to 78 families. Among the plant familes Fabaceae, Asteraceae,, Acanthaceae and Lamiaceae are dominant families, comprising higher number of bee forage species composition in the area. These families are also reported to be the most species-rich families in the Flora of Ethiopia and Eritrea [10], [1]. Moreover the Fabaceae is one of the pollinator dependent plant family and contributing for honey production and serving as food source for other pollinators. Analysis of life forms of the plant revealed that the herbaceous flora contributes for 35% of the total species composition of the area. This could be due to removal of shrubs and trees from farmland for cultivation and forest encroachment favoring the growth of herbaceous flora. This is in agreement with [11] agricultural practices favor for the growth various weeds and other herbaceous flora.

Based on plant inventory and honey pollen analysis indicated that the bee forage diversity of bee plants was not vary among the districts but species richness was highest for Gomma and Omonda. This is due to the area located in one of the Biodiversity hot spots which is known as Eastern Africa Biodiversity hot spots which is known for high plant diversity. Similar finding by [12] in the Borena Zone of Oromia, Ethiopia. The PCA analysis of honey samples of the study districts revealed that, *Guizotia spp*, contributed more than 72% of the pollen count.

### Identification of major bee forages from pollen trap

The identification of bee-collected pollen loads from pollen traps indicated that not only honeybees collected pollen from plant species but also it shows the relative importance of each plant species as source of pollen for honeybees. A total of 41 plant species were identified from pollen trap, however the large quantity of pollen came from only a few plan species such as *Bidens spp, Eucalyptus spp, Guizotia scabra, Plantago lanceolata, Masea lanceolata, Trifolium* spp, *Vernonia spp, Terminalia schimperiana* and *Zea mays*. The contribution of each species as major source of pollen is due to abundance and potentiality of plant for pollen. This result is also supported by findings of [8] and [13], who reported that honeybees forage on more productive and profitable plants that provide necessary nutrition nearby their hives.

### Seasonal pollen collection

The highest fresh weight of pollen was collected during the months of October, November and December in the area due to extended rainy season resulting for longer duration of plants in flower. In addition, there are some drought tolerant species such as *Vernonia amgydalina, Bersama abyssinica, Terminalia schimperiana* and *Coffea arabica*. This is agreeing with [14], [15], [16] who reported that *Vernonia Spp, Coffea arabica, and Croton macrostachyus*, are the major bee flora flowering both in dry period and small rainy season in the Gomma district. According to the same author *Vernonia amygdalina* is a very valuable honey source plant in the area. In this district low pollen collection was occurred during the months of March due to availability of low moisture in the soil resulting drying of flowers. Relatively higher pollen collection was recorded during the months of April, May and June for Gomma, Tolay and Omonada districts due to availability of *Croton macrosatchyus* and similar finding also reported by [14], [15].

### Flowering calendar of the bee forages in west and south west shoa

From field observation and pollen load collection the highest percentage of flowering bee forages occurred during September to November reaching peak flowering in October in all study districts. The availability of higher proportion of flowering species during October to November is due to occurrence of summer rains in June, July and August and at the end of this rainy period, majority of the plants come into flower. Similar flowering pattern was observed for Tolay, Gomma and Omonada in which majority of bee forages flowering from October to December due to extension of summer rainfall creating availability of enough moisture until end of December. Exceptionally some plant species including *Vernonia amygadlina*, and *Coffea arabica* abundantly flowering from January to February and minor honey flow is expected in Gomma district. This study also agreement with [13], [8] indicated that *Vernonia Spp*., *Coffea arabica, Schefflera abyssinica* and *Croton macrostachyus were* common honey bee floras in Jimma zone flowering from January to May. As reported [18] small honey harvesting periods was identified in Gomma district which depend on the type of flowering plants and rainfall patterns.

### Pollen analysis of honey

Pollen spectra of honey analysis revealed that a variety of pollen source plants were identified ranging from predominate, secondary pollen and minor important. The degree of the dominance of the bee forages depend on abundance, nectar and pollen potential of the plants. Based on honey pollen analysis the highest bee forage diversity of bee plants was found in the honey sample collected from Tolay and Ejere as compared to the rest of the districts since the study area is located in central highland which is known for high plant diversity. Similar finding by [12]in the Borena Zone of Oromia, Ethiopia. The PCA analysis of honey samples of the study districts revealed that, *Guizotia spp*, contributed more than 72% of the pollen count. The dominance of *Guizotia* pollen from honey can be attributed to widespread distribution in agricultural lands, fallow land and since it is associated weed of major crops that grow within and at the margin of field crops. This is agreeing with [13],[19],[20] who reported that *Guizotia scabra* was a predominant monofloral honey source plant in east and west shoa and Borena zones of Oromia. The availability of coffee honey and croton honey in Jimma zone were reported by[13] and [17].

### Conclusion and Recommendation

Based on direct field observation, and pollen load collection 103 plant species were identified and herbaceous flora comprising the major portion of bee forages in the area. The higher proportion of the pollen weight was collected from a few plant species including *Guizoti*a spp, *Vernonia* spp, *Eucalyptus camaludensis, Echinops macrostchyus,s Bidens pilosa, Masea lanceolata, Planatago lanceolata, Echinopes macrostachus*. Majority of plant species flower two times a year during September to November and April to May in most districts. Based on floral calendar, the scarcity of pollen collection was occurred in early march and July and feeding colonies may require. Thus creation of awareness and on farm demonstration of the important of floral calendar for appropriate management of honeybee colonies is recommended to increase frequency of honey harvest. Moreover *Masea lanceolata* was identified as major bee forages during rainy period and further study on the rapid propagation methods is recommended.

## Acknowledgment

We acknowledge the Holeta Bee Research Center and Oromia Agricultural Research Institute for providing required facilities and logistics. Our sincere thanks also extended to Ms. Konjit Asfaw, Berhane Tadesse and Mr Tesfaye Abera and our driver Bekele Gemechu for helping us during field data collection.

## Appendix1. Checklist of plant species from Jimma zone

| Plant species | Habit | Family | Flowering period | Food source |
| --- | --- | --- | --- | --- |
| *Acacia abyssinica* | Tree | Fabaceae | Sept- Nov | N,P |
| *Acacia seyal* | Tree | Fabaceae | Sept- Nov | N,P |
| *Acacia tortoils* | Tree | Fabaceae | April- May | N,P |
| *Acanthus senni* | Shrub | Acanthaceae | Sept-Oct | N |
| *Achranthes aspera* | Herb | Amaranthaceae | Sept-Oct | P |
| *Agava sisalina* | Shrub | Agvaceae | Dec-Jan | N |
| *Ageratum houstonianum* | Herb | Asteraceae | Sept- Dec | P |
| *Albizia gummifera* | Tree | Fabaceae | Sept-Oct | N,P |
| *Bersama abyssinicum* | Tree | Meliaceae | Sept-Dec | N,P |
| *Bidens pillosa* | Herb | Asteraceae | Oct- Dec | P |
| *Bidens prestinaria* | Herb | Asteraceae | Sept- Oct | P |
| *Brassica spp* | Herb | Brassicaceae | Sept- Oct | N,P |
| *Brucea antidysenterica* | Shrub | Simaroubaceae | Sept- Oct | P |
| *Caesalpinia decapetala* | Climber | Fabaceae | Oct- Dec | N,P |
| *Calpurnia aurea* | Shrub | Fabaceae | Oct- Dec | P |
| *Capiscum annum* | Herb | Solanaceae | Sept- Oct | P |
| *Carica papya* | Shrub | Caricaceae | Sept- Dec | N,P |
| *Carrisa edulis* | Shrub | Apocynaceae | Sept- Dec | P,N |
| *Catha edulis* | Shrub | Celastraceae | Sept- Nov | P |
| *Caylusea abyssinica* | Herb | Resedaceae | Oct- Dec | N |
| *Celtis africana* | Shrub | Ulmaceae | Sept-Nov | P |
| *Cicer arietinum* | Herb | Fabaceae | Dec-Jan | N,P |
| *Citrus sinensis* | Shrub | Rutaceae | Sept-Nov | N,P |
| *Clausena anisata* | Shrub | Rutaceae | Sept-Nov | P,N |
| *Climatis hirsuta* | Climber | Ranunculaceae | Nov-Dec | P,N |
| *Coffea arabica* | Shrub | Rutaceae | Jan- Feb | N,P |
| *Cordia africana* | Tree | Boraginaceae | Sept- Oct | N,P |
| *Croton macrostachyus* | Tree | Euphorbiaceae | May- June | N,P |
| *Cynotis barbarta* | Herb | Commelinaceae | Sept-Oct | P |
| *Cytisus proliferus* | Shrub | Fabaceae | July- Aug | N,P |
| *Datura arborea* | Shrub | Solanaceae | Sept- Jan | N,P |
| *Datura stramonium* | Shrub | solanaceae | Sept- Nov | N,P |
| *Desmodium unicilayum* | Herb | Fabaceae | Sept- Oct | N,P |
| *Dombeya torrida* | Tree | Stericulaceae | Sept- Oct | N,P |
| *Dovalis caffra* | Shrub | Flocourtiaceae | Mar- May | P |
| *Echinopes spp* | Herb | Asteraceae | Sept-Oct | P |
| *Ehretia cymosa* | Shrub | Boraginaceae | Sept-Oct | N,P |
| *Ekbergia capensis* | Tree | Meliaceae | Sept-Oct | N,P |
| *Erhrina abyssinica* | Tree | Fabaceae | Sept- Oct | N |
| *Eucalyptus camadulensis* | Tree | Myrtaceae | Mar- April | N,P |
| *Eucalyptus globulus* | Tree | Myrtaceae | Mar- April | N,P |
| *Euclea schimperi* | Tree | Ebenaceae | Sept-Oct | N,P |
| *Galinosoga parviflora* | Herb | Asteraceae | Sept-Oct | P |
| *Glycine weightii* | Herb | Fabaceae | Sept-Oct | P |
| *Terminalia brownii* | Tree | Combretaceae | Dec-Jan | N,P |
| *Grvillea robusta* | Tree | Proteaceae | Sept-Nov | N |
| *Polyscias fluva* | Tree | Araliaceae | Sept-Nov | N,P |
| *Guizotia scabra* | Herb | Asteraceae | Oct-Nov | P |
| *Haygrophilia auriculata* | Herb | Acanthaceae | Sept-Nov | N,P |
| *Hypoestes triflora* | Herb | Acanthaceae | Sept-Oct | N,P |
| *Impomea spp* | Herb | Impomeaceae | Sept-Oct | N,P |
| *Jacarnda mimosifolia* | Tree | Bignoniaceae | Sept-Oct | N,P |
| *Justicia bizunesh* | Shrub | Acanthaceae | Sept-Oct | N,P |
| *Justicia schimperana* | Shrub | Acanthaceae | Sept-Dec | N,P |
| *Casimiroa edulis* | Shrub | Rutaceae | Sept-Dec | N,P |
| *Leucas marienthesis* | Herb | Lamiaceae | Sept-Dec | N,P |
| *Leucaena leucocephala* | Tree | Fabaceae | Sept-Dec | N,P |
| *Lippia adoensis* | shrub | Verbenaceae | Sept-Oct | N,P |
| *Malus silvestris* | shrub | Roseaceae | Sept-Oct | N,P |
| *Mangifera indica* | shrub | Anacardiaceae | Sept-Oct | N,P |
| *Masea lanceolata* | Shrub | Myrisenaceae | Aug- sept | P |
| *Mayntus obscura* | Shrub | Calestraceae | Sept-Oct | P |
| *Maytenus arbutifolia* | shrub | Calestraceae | Sept-Oct | P |
| *Morings stenopetala* | Tree | Moringaceae | Sept-Oct | N,P |
| *Morus alba* | shrub | Moraceae | Sept-Oct | P |
| *Musa paradisica* | Shrub | Musaceae | Sept-Oct | N,P |
| *Myrica salicifolia* | Shrub | Myricaceae | Sept-Oct | N,P |
| *Nicandra physaloides* | Herb | solanaceae | Sept-Oct | N,P |
| *Ocimum lamifolium* | Herb | Lamiaceae | Oct -Dec | N,P |
| *Ocimum urticifolium* | Herb | Lamiaceae | Sept-Oct | N,P |
| *Octea kenysis* | Shrub | Rutaceae | Sept-Oct | N,P |
| *Olea capensis* | Shrub | Oleaceae | May | N,P |
| *Opuntia ficus indica* | Shrub | Euphorbiaceae | Sept-Oct | N,P |
| *Pavonia urens* | Herb | Malvaceae | Sept-Oct | N,P |
| *Persea americana* | Tree | Luraceae | Sept-Oct | N,P |
| *Phonix reclinata* | Tree | Areceae | Sept-Oct | N,P |
| *Phytolacca dodocandra* | Shrub | Phytolaceae | Sept- Oct | N,P |
| *Piper capense* | Shrub | Piperaceae | Sept- Oct | P |
| *Plantago lanceolata* | Herb | Plantaginaceae | Sept-Jan | P |
| *Plectranthus punctata* | Herb | Lamiaceae | Sept-Oct | N |
| *Premena schimeprina* | Shrub | Premiaceae | Oct-Nov | N,P |
| *Psidium guajava* | Shrub | Myrtaceae | Sept-Oct | N,P |
| *Capparis tomentosa* | Climber | Cappridaceae | Sept-Oct | N,P |
| *Rhamnus prinoides* | Shrub | Rhamnaceae | Sept-Oct | P |
| *Ricinus communis* | Shrub | Euphorbiaceae | Sept-Oct | N,P |
| *Rubus steudneri* | Shrub | Roseaceae | Sept-Dec | N |
| *Rumex nervosus* | Shrub | Polygonaceae | Sept- Dec | P |
| *Schefflera abyssinica* | Tree | Araliaceae | March- April | N |
| *Sida rhombifolia* | Shrub | Malvaceae | Sept- Dec | P |
| *Sorghum bicolor* | Herb | Poaceae | Sept-Oct | P |
| *Stereospermum kunthianum* | Tree | Bignoniaceae | Sept-Oct | N,P |
| *Syzygium guineense* | Tree | Myrtaceae | Feb-Mar | N,P |
| *Tagetes minuta* | Herb | Asteraceae | Sept-Oct | N,.P |
| *Terminalia schimperiana* | Tree | Combretaceae | Dec-Jan | P |
| *Tobacco tubersum* | Herb | solanaceae | Sept-Nov | N |
| *Urerea hypodenderon* | Climber | Urticaceae | Sept-Nov | P |
| *Vernonia amygdalina* | Shrub | Asteraceae | Dec- Jan | N,P |
| *Vernonia auriculifera* | Shrub | Asteraceae | Dec- Jan | N,P |
| *Vernonia leopoldii* | Shrub | Asteraceae | Nov-Dec | N,P |
| *Vernonia thomsonii* | Shrub | Asteraceae | Dec- Jan | N,P |
| *Zea mays* | Herb | Poaceae | Sept | P |
| *Zehneria scabra* | Herb | Cucurbitaceae | Sept-Nov | N |

N= nectar, P=pollen, NP= Nectar and Pollen

## References

[1] Admassu A. Wakjira, Kibebew W. Amssalu B.. &, Ensernu K. 2014. Honey bee forages of Ethiopia. United Printers

[2] Haftom K. and Samuel G. 2016. Floral calendar establishment of Major honeyplants in northwestern zone of Tigray, Ethiopia. International Journal of Scientific & Engineering Research, 7,(9)

[3] Nigel and Anna 2019. Climate effects on the onset of flowering in the United Kingdom. Fox and Jönsson Environ Sci Eur. 31(89) pp 3–13.

[4] Fabian N., Stephan H. and Ingolf S. 2018. The influence of temperature and photoperiod on the timing of brood onset in hibernating honey bee colonies

[5] Admassu, A. 2003. Botanical Inventory and Phenology in Relation to Foraging Behavior of the Cape Honeybees (Apis mellifera capensis) at a Site in the Eastern Cape (MSc thesis). Rhodes University, South Africa

[6] Admassu A. & P. Phillipson R. Hepburn 2006. Floral resources of Apis mellifera capensis in the fynbos biome in the Eastern Cape, south Africa. J. of African Entomology 14(1)

[7] Abebe, J., Amssalu, B., & Kefelegn, K. 2014. Floral phenology and pollen potential of honey bee plants in North East dry land areas of Amhara region, Ethiopia. IOSR Journal of Agriculture and Veterinary Science: 7(5): 36–49.

[8] Tolera K. and Dajene T. 2014. Assessment of the Effect of Seasonal Honeybee Management on Honey Production of Ethiopian Honeybee (Apis mellifera) in Modern Beekeeping in Jimma Zone

[9] Louveaux, J., Maurizio, A., & Vorwohl, G. 1978. Methods of Melissopalynology. Bee World, 51(3), 125–138.

[10] Ensermu K. and Sebsebe D. 2014. Diversity of vascular plant taxa of the flora of Ethiopia and Eritrea. Ethiop. J. Biol. Sci. 13: 37–45.

[11] Admassu A. Ensermu K., Teshome S., Peter G.P, Lulsegde B., Campos 2017: Proximate composition and antioxidant power of bee collected pollen from moist Afromontan forests in southwest Ethiopia Vol. 7(3): 83–95.

[12] Tura, B., & Admassu, A. 2018. Pollen analysis of honey from borana zone of Oromia southern Ethiopia J. Apic. 63(2)[13]

[13] Debissa L, Admassu A. 2009. Identification and Evaluation of Bee Flora resources in arid and Semiarid Agro-ecological Zones of South east of Oromia. In proceedings of 17th Annual Conference of Ethiopia society of Animal production, Addis Ababa, Ethiopia.

[14] Abera H. 2017. Identification of Honey Source Bee Floras during Major and minor Honey harvesting Seasons in Jimma Zone, Southwest Ethiopia. Journal of Environment and Earth Science 7(3): 25–32

[15] Chala K., Taye E., Kebede D. Tadesse T. 2012.. Opportunities and challenges of honey Production in Gomma district of Jimma zone, South-west Ethiopia.

[16] Mohammed A., Kebede D. Zemene W. 2016. Assessment of Beekeeping Practices in Shabe and Seka Chekorsa Districts of Jimma Zone, Southwestern Ethiopia. European Journal of Biological Sciences 8 (2): 45–55

[17] Tura B. and Admassu A. 2020. Bee flora diversity in different vegetation communities of Gesha-Sayilem forest in Kaffa Zone, south-western Ethiopia. Plants and Environment 2(4): 138–148

[18] Nuru, A., (2007). Atlas of pollen grains of major honeybee flora of Ethiopia. Ethiopian Society of Animal production, Addis Ababa, Ethiopia, p. 152.

[19] Mesert G. (2018). Physicochemical and Antioxidant Properties of Honey and Pollen from West Shoa, Ethiopia Msc. Thesis, Addis Ababa University

[20] Tura B. and Admassu A., (2019). Bee flora resources and honey production calendar of Gera Forest in Ethiopia Asian Journal of Forestry 3(2):69–74

